# Defying expectations: Extended lifespan and limited age-related telomere shortening in the tropical bat species, *Molossus molossus*

**DOI:** 10.1101/2025.10.30.685550

**Authors:** Tadhg Lonergan, Megan L. Power, Luisa F. Gomez, Sebastien Riquier, Igor Sukhikh, Melissa Lopez, Rachel Page, Frederic Touzalin, Dina K. N. Dechmann, Emma C. Teeling

**Affiliations:** School of Biology and Environmental Science, University College Dublin, Belfield, Dublin 4, Ireland; School of Biological Sciences, University of Bristol, Life Sciences Building, 24 Tyndall Avenue, Bristol BS8 1TQ, UK; Universidad de Costa Rica, Sede Rodrigo Facio, Golfito, 60701, Costa Rica; Smithsonian Tropical Research Institute, Luis Clement Avenue, Bldg. 401 Tupper, Ancon, Panama 0843-03092, Republic of Panama; Max-Planck Institute for Animal Behavior; Radolfzell, 78315, Germany; University of Konstanz; Konstanz 78464, Germany

**Keywords:** telomeres, chiroptera, longevity, short-lived, mark-recapture

## Abstract

Telomeres are key biomarkers of cellular ageing, yet their dynamics remain poorly studied in tropical and short-lived bat species. Here, we present the first investigation of telomere length across age in *Molossus molossus*, a tropical bat historically categorised as the shortest-lived bat on record. Through a multi-year mark-recapture study in Gamboa, Panama, we sampled 492 individuals (*n* = 317 females, 175 males) and documented a female *M. molossus* surviving to at least 13 years of age, more than doubling the previously reported maximum lifespan of 5.6 years. Across the population, relative telomere length (rTL) showed no overall significant decline with age. No evidence was found for sex-specific rates of telomere attrition. Rather these results suggest that males and females follow parallel age-related telomere trajectories, with any sex differences primarily reflecting differences in mean telomere length rather than ageing dynamics. Overall, the findings here challenge previous assumptions about the lifespan and ageing biology of *M. molossus*. They demonstrate that telomere maintenance is not limited to temperate bats, show that sex differences in telomere biology are subtle and species-specific, and reinforce the value of long-term field studies for understanding ageing processes in the wild.

## Introduction

Vertebrate lifespan shows extraordinary variation, from species such as pygmy goby (*Eviota sigillata*) living ∼8 weeks (Depczynski and Bellwood, 2005) to the Greenland shark (*Somniosus microcephalus*) living up to 500 years (Nielsen et al., 2016). Body size reflects much of this range, with large animals typically living longer than smaller animals due to lower extrinsic mortality and slower metabolic turnover (Speakman, 2005). Within mammals, in particular, there is a log-linear relationship between body mass and maximum lifespan, where larger species tend to live longer than smaller species (Austad, 2010). However, bats (order Chiroptera) defy this relationship, displaying exceptional longevity and are the longest-lived mammals relative to their body size (Austad, 2010, Foley et al., 2020). Many bat species can live considerably longer than terrestrial nonflying mammals of a similar size (Healy et al., 2014, Seim et al., 2013, Wilkinson and Adams, 2019). Indeed, some small bodied bats exhibit exceptional longevity. For example, *Myotis brandtii*, recaptured at over 41 years of age despite weighing only ∼7 g, exhibits a longevity quotient (LQ; the ratio of observed lifespan to that predicted from body mass) of 8.23, living more than eight times longer than expected for a mammal of its size (Foley et al., 2018; Podlutsky et al., 2005). This remarkable lifespan has positioned bats as valuable models for ageing research (Cooper et al., 2024, Teeling et al., 2018).

One promising approach for investigating ageing in wild mammals is telomere dynamics, a widely used biomarker of biological ageing across taxa (Lopez-Otin et al., 2023). Telomeres are short repetitive nucleotide sequences which protect the ends of linear chromosomes and prevent genomic instability (Blackburn, 1991). In most organisms, telomeres shorten with each round of cell division due to incomplete end replication during cell division, known as the “end replication problem” (Levy et al., 1992, Watson, 1972). Telomere length can also shorten when exposed to sources of cellular damage, such as reactive oxygen species (Armstrong and Boonekamp, 2023, Reichert and Stier, 2017, von Zglinicki, 2002). When telomere length becomes critically short, replicative cell senescence is triggered, halting cell division (Harley et al., 1990, Jones-Weinert et al., 2024). In some species, the enzyme telomerase can restore telomere length, counteracting this shortening, although in most mammals telomerase activity is downregulated in somatic tissues after development. This is thought to have evolved as a tumour-suppressive mechanism, reducing the risk of malignant transformation at the cost of progressive cellular senescence (Gomes et al., 2011, Shay and Wright, 2019). However, studies focusing on long-lived bat species, particularly within the *Myotis* genus, have revealed unique telomere dynamics.

Unlike most mammals, where telomeres shorten with age, in the genus of bats with the longest maximum lifespan relative to body size, *Myotis,* there is no correlation between telomere length and chronological age (Foley et al., 2018, Foley et al., 2020, Ineson et al., 2020). This occurs without detectable telomerase activity in somatic tissues, suggesting alternative mechanisms for preserving telomere integrity (Foley et al., 2018, Foley et al., 2020). However, little is known about telomere dynamics across bats. Despite more than 1500 bat species recorded globally (Simmons & Cirranello, 2025), nearly all studies to date are based on longer-lived temperate insectivorous bats (Foley et al., 2018, Foley et al., 2020, Ineson et al., 2020, Power et al., 2022, Power et al., 2023). A recent study in the tropical bat *Phyllostomus hastatus* provided the first evidence of age-related telomere shortening in a non-temperate species, based on cross-sectional sampling and methylation-derived age estimates (Rayner et al., 2025). However, *P. hastatus* is a relatively long-lived tropical bat (females live up to 22 years; Wilkinson and Adams, 2019) and therefore a gap remains in research concerning telomere dynamics in short-lived bat species, with no known studies to date. Investigating such species could clarify whether short-lived bats follow more conventional mammalian telomere ageing patterns, and whether telomere maintenance is restricted to certain long-lived temperate lineages (e.g., *Myotis*).

*Molossus molossus,* commonly known as Pallas’s mastiff bat or the velvety free-tailed bat, offers the ideal short-lived contrast. Based on reliable data, *M. molossus* has been categorised as the shortest-lived bat species on record (Tacutu et al., 2018), with a maximum recorded lifespan of about 5.6 years in the wild and median survival for wild females estimated to be approximately 1.8 years (Gager et al., 2016a). *M. molossus* is among the most widespread and abundant species in the Neotropics, with a distribution ranging from southern Mexico through Central and South America and the Caribbean (Loureiro et al., 2021, Reid, 2006, Simmons, 2005). In Panama, stable roosting groups dominated by adult females have been well studied (Dechmann et al., 2010, Gager et al., 2016a). These groups are typically joined by one or two adult males with frequent replacement of harem males. *M. molossus* is strictly tropical and does not hibernate, however they reduce heart rate, along with metabolic rate by >70% while maintaining body temperature around 32 °C, similar to the daytime torpor employed by temperate zone bats (O’Mara et al., 2017, Wojciechowski et al., 2007). Such a trait may reduce oxidative damage and influence telomere dynamics.

Here, we provide the first assessment of telomere length dynamics in *Molossus molossus* leveraging over 15 years of mark-recapture data from wild populations in Panama. Utilizing population-level qPCR estimates, we investigated age-related telomere changes in this tropical insectivorous bat. Given its previously reported short lifespan, we hypothesised that telomere length would decline with advancing age, in this putative short-lived bat.

## Materials and Methods

### Study population

This study focuses on a long-term mark-recapture programme for *Molossus molossus* in Gamboa (N 09,07; W 079,41) Panama, initiated in 2008 (Dechmann et al., 2010, Gager et al., 2016a). Gamboa is located near the banks of the Panama Canal and the Chagres River, surrounded by Soberanía National Park, and the study area is covered by semi-deciduous tropical lowland rainforest (Leigh, 1999, Windsor, 1990). Between 2008 and 2014, 14 social groups were captured on 81 occasions with approximately 500 individuals PIT-tagged, producing the first survival and longevity estimates for this species (maximum lifespan 5.6 years; Gager et al. 2016). Smaller yearly mark-recapture catches have continued since 2014, up until the start of this study.

Individuals used in this study were caught and sampled over the course of four years (2021 – 2024). Sampling was carried out consistently over two-three weeks in October/ November of each year, where the majority of female bats have finished lactating. In addition to these annual samplings, a subset of individuals (n = 109) were also opportunistically sampled during the summer months (May - August) in 2022 and 2023. All capture and handling of bats was carried out with appropriate permission from the Autoridad Nacional del Ambiente in Panama and with approval from the Institutional Animal Care and Use Committee of the Smithsonian Tropical Research Institute.

### Capture and sampling

In total, 30 sampling events were conducted across 14 different roost sites in Gamboa, (n = 583 capture events). Roosts were selected based on prior knowledge of *Molossus* bat presence and through systematic searches in human-made structures, typically crevices on the exterior of buildings. Bats were caught using mist nets placed at roost entrances, shortly after sunset. Upon capture, all bats were placed in soft cloth bags prior to processing. Bats were sexed, weighed (to an accuracy of ± 0.1g using a precision weight balance), and forearm length was measured (to an accuracy of ± 0.1mm using Vernier callipers). Newly caught individuals were marked with a subcutaneous Passive Integrated Transponder (PIT) tag (2.12x11mm, 0.1g, Trovan® & Biomark™), containing a unique 10-digit code which can be read using passive Radio-Frequency Identification (RFID) readers and facilitate the identification of individuals at each subsequent recapture. Wing punches were taken from all individuals at every capture using a sterile 3mm biopsy punch and immediately flash frozen in the field in liquid nitrogen (LN_2_).

Blood samples were also collected for each bat, by piercing the cephalic vein along the forearm with a sterile needle (27 gauge), creating a drop of blood on the patagium surface from which a volume of <100 μL was pipetted into a cryotube (2 ml, Nalgene labware). Haemostatic gel was applied on the vein to prevent further bleeding and all bats were checked before release to ensure bleeding had completely stopped (Huang et al., 2016). Blood samples were also immediately flash frozen in LN_2_. All bats were released in the same area in which they were captured. As *M. molossus* bats are unable to take flight from low levels, all bats were released onto nearby trees where they could climb to an appropriate height before taking flight. Bats were monitored post-release to ensure they successfully took flight.

### Age determination

The age of the bat was calculated based on the first time the bat was caught and PIT tagged. Captures were timed to occur around the time juveniles were fledged but still present in the maternal roost. If a bat was caught and tagged as a juvenile for the first time, its age was recorded as 0 and it was considered known aged. If a bat was caught for the first time as an adult, then it was assigned the age of “1+” indicating they were at least one year old at time of capture. If re-captured a year later this individual was recorded as “2+” and so forth. Juvenile bats were differentiated from adults based on the degree of ossification of the epiphyseal cartilage in the metacarpal-phalangeal joints (Kunz and Anthony, 1982) as well as by darker pelage colour than adults.

### Species identification

While this study was designed to focus on *Molossus molossus*, the cryptic species *Molossus coibensis* was identified at a subset of roosts. Morphological classification was performed using forearm length, body mass, and pelage colour (Gager et al., 2016b). To validate the field morphological assignments, we confirmed the species identity of a subset of individuals (n = 72) using molecular phylogenetics. Total RNA was extracted from whole blood using an RNAzol BD kit, following the manufacturer’s instructions with minor modifications. The full RNA extraction from bat blood protocol is described in detail by Huang et al., (2016). Quantity and quality of RNA were assessed using a Bioanalyzer 2100 (Agilent Technologies). All samples that met the criteria of having >2 μg of total RNA and an RNA Integrity Number score >8.0 were chosen for RNA-Seq Illumina library preparation. The generated RNAseq data were quality trimmed using *Trim Galore!* v0.6.4 (Martin, 2011), then mapped on the *M. molossus* genome from Jebb et al. (2020) with STAR software v2.7.10b_alpha using default parameters (Dobin et al., 2013). Genomic variants were extracted from expressed regions using *bcftools* “mpileup” and “call” functions (Danecek et al., 2021), and filtered for a minimum quality of 20 and coverage of 10. The BRCA1 sequence for each individual was reconstructed by applying the filtered variants to the longest annotated isoform of the *M. molossus* reference, BRCA1_x2, using *bcftools* consensus. A *Myotis myotis* BRCA1 sequence was used as an outgroup (Jebb et al., 2020). Multiple sequence alignment was performed using the MUSCLEv3 program (Edgar, 2004) with default parameters. A maximum likelihood phylogeny was constructed using IQ-tree (Trifinopoulos et al., 2016), with the substitution model GTR+G+I determined by the software based on the lowest AICc (Corrected Akaike Information Criterion) score. Branch support was assessed using two statistical tests available in IQ-tree: SH-like aLRT (approximate Likelihood Ratio Test, 1,000 replicates) and UfBoot (ultrafast bootstrap, 1,000 replicates).

### DNA extractions and qPCR assay

DNA was extracted from wing biopsies using the membrane filter-based extraction kit - Promega Wizard SV DNA Extraction Kit (catalogue no. A2371, Promega Corporation, Madison, WI). This 96 well extraction protocol was partially automated using the Hamilton Star Deck liquid Handling Robot but otherwise followed the manufacturer’s instructions. DNA concentration and purity was quantified using a Nano Drop 8000 Spectrophotometer (ThermoScientific). Wing tissue has been shown to be a suitable proxy for global tissue telomere dynamics in bats (Power et al., 2021). Relative telomere length (rTL) was measured by real-time quantitative PCR (qPCR) as the concentration of telomeric DNA relative to a single copy gene (SCG) that is constant in number and optimised for use in bats. qPCR amplifications were carried out using the Applied Biosciences Quantstudio 7 Flex Real-Time PCR. qPCR assay followed a modified version of the Cawthon (2002) protocol adapted for bats as described in Foley et al. (2018). Samples were run on either 96-well or 384-well plates, with six technical replicates. A “golden sample” calibrator prepared from pooled *M. molossus* wing tissue was run in triplicate on each plate to account for among-plate variation as well as a negative control to detect contamination.

Raw data was exported from the qPCR machine and the software package LinRegPCR (Ruijter et al., 2009) was used to calculate baseline correction, per well amplification efficiencies and Cq values (the number of cycles the qPCR amplification curve crosses a set fluorescence threshold). Data were normalised using the ‘golden sample’ as reference across both telomere and SCG reactions. The average Cq was calculated across the six replicates for each sample for both amplicons. rTL was calculated for each sample, following Pfaffl (2001). Co-efficient of variation (CoV) thresholds were applied: 5% for telomere reactions and 2.5% for single-copy gene (SCG) reactions (Foley et al., 2018, Foley et al., 2020).

Average reaction efficiencies (mean ± SE) across the 12 qPCR plates were 1.984 ± 0.012 for the SCG reaction and 1.931 ± 0.026 for the telomere reaction. Intra-plate repeatability (intraclass correlation coefficient, ICC) calculated with the rptR package (Stoffel et al., 2017) was 0.971 ± 0.002 (95 % CI [0.967–0.974], n = 3528 technical replicates, P < 0.001) for SCG and 0.957 ± 0.003 (95 % CI [0.952–0.962], n = 3528 technical replicates, P < 0.001) for telomere. (1,000 parametric bootstraps; 1,000 permutations, rptR). Inter-plate consistency was calculated on the subset of samples analysed in triplicate on two separate plates. Plate-to-plate effects accounted for only ≈ 6 % of the telomere Cq variance (ICC_plate = 0.067 ± 0.070) and ≈ 16 % of the SCG variance (ICC_plate = 0.157 ± 0.111). However, differences among samples remained the dominant source of variation (ICC_sample = 0.823 ± 0.065 for telomere; 0.652 ± 0.091 for SCG; all P < 0.001).

### Statistical analysis

All statistical analyses were carried out in R v.3.6.3 (R Core Team, 2021). As residual diagnostics indicated deviations from normality and heteroscedasticity, a Box-Cox transformation (Box and Cox, 1964) was applied to the response variable to improve model fit. rTL measures were subsequently transformed to z-scores to facilitate comparability across studies (Verhulst, 2020). Linear mixed effect models were constructed using the R packages MASS (Venables, 2002) and lme4 (Bates et al., 2015) to investigate relationships between rTL, age and sex. Models included random effects for individual ID, qPCR assay plate, site, and sampling year to control for repeated measures and non-independence in the data. The best-fitting model was selected based on Akaike’s Information Criterion corrected for small sample sizes (AICc), and residuals were examined to ensure model assumptions were met. We also tested for random effects of capture month, but its inclusion did not improve model fit or affect fixed effect estimates and was therefore excluded from the final models. To test for sex-specific age-related telomere dynamics, a sex × age interaction term was included.

Given the presence of individuals at extreme ages relative to the overall distribution (see Results below), we fitted robust linear mixed effects models using the robustlmm package (Koller, 2016). Robust linear mixed modelling differentially weights residuals to reduce the influence of outliers, helping to address departures from model assumptions and limited sample sizes while retaining all observations (Filzmoser and Nordhausen, 2021). Robust models were fitted to both the full dataset and the dataset excluding the oldest individual as part of a sensitivity analysis to evaluate the influence of extreme-age observations on parameter estimates. Robust mixed models were fitted with the fixed effects of age and sex and we additionally tested a model including a sex × age interaction. As rlmer does not report p-values, we assessed statistical significance using 95% parametric bootstrap confidence intervals (10000 iterations) implemented in the confintROB package (Mason et al., 2024). Fixed effects were considered significant when the confidence interval did not include zero.

In our dataset, females were more numerous than males due to the harem based structure of most *Molossus* colonies. To account for the resulting imbalance in sex and age distributions, we applied an iterative subsampling approach to test the robustness of sex differences in rTL when age distributions were equalized (Rayner et al., 2025). For each of 10,000 iterations, males were randomly matched to females of the same exact age, and a simple linear model was used to test the effect of sex on rTL. A Wilcoxon rank-sum test was used in each iteration to confirm matched age distributions between sexes. The proportion of iterations where the sex effect was statistically significant (*p* < 0.05) was recorded. Additionally, the distribution of sex effect estimates across all iterations was plotted to assess the consistency and direction of any observed sex bias.

The *M. molossus* dataset contains many individuals which were first captured as adults and thus have an approximate age e.g. ‘1+’. As such the data was analysed two ways: using individuals of known age (n = 151) and in a minimum age based analysis (n = 466) where the ‘x+’ ages were converted to whole ages and treated as the minimum age of the bat, to maximise usable data.

## Results

### Capture and species identification

Over the course of this study, 583 captures (428 individuals) of the genus *Molossus* were recorded across 14 different roosts sites in Gamboa, Panama from 2021 to 2024. Species identification was carried out on all individuals based on morphological characteristics and, on a subset (n = 72) using BRCA1 gene transcripts derived from Illumina blood-transcriptome reads. 492 bats were morphologically identified as *M. molossus* and 91 as *M. coibensis.* All genetically identified individuals matched their infield, morphological classification, confirming the reliability of morphological criteria used in the larger dataset. A morphological comparison between species shows clear species groupings based on forearm length and body mass (Figure S1). *M. coibensis* individuals were generally smaller in forearm length but heavier in body mass relative to *M. molossus*. Phylogenetic reconstruction based on *BRCA1* sequences revealed distinct species-level clustering consistent with field identifications for *M. molossus* and *M. coibensis* (Figure S1). This analysis confirmed the placement of each specimen within the expected *M. molossus* or *M. coibensis* clades. Of the 492 *M. molossus* captures, 367 unique individuals were identified, of which 317 were females and 175 males. Among these, 163 were of known age (tagged as juveniles), while the remaining 329 were assigned a minimum estimated age. A total of 82 individuals were recaptured at least once, giving a recapture rate of 22.3%. Capture and recapture data for *M. molossus* are summarised in Table S1.

### *Molossus molossus* - New longevity record

A significant finding of this study was the identification of an individual that was at least 13 years old at her last capture in 2024, representing the oldest known *M. molossus* recorded to date (Figure 1). Bat 0006C8684C was originally caught from the same roost and tagged as an adult in 2012. Previously, the maximum reported lifespan for the species was 5.6 years and this new estimate has more than doubled the recorded maximum longevity for this species. This new age record also changes the longevity quotient of *M. molossus* from 0.99 to 2.37 (Figure 1).

**Figure 1:**
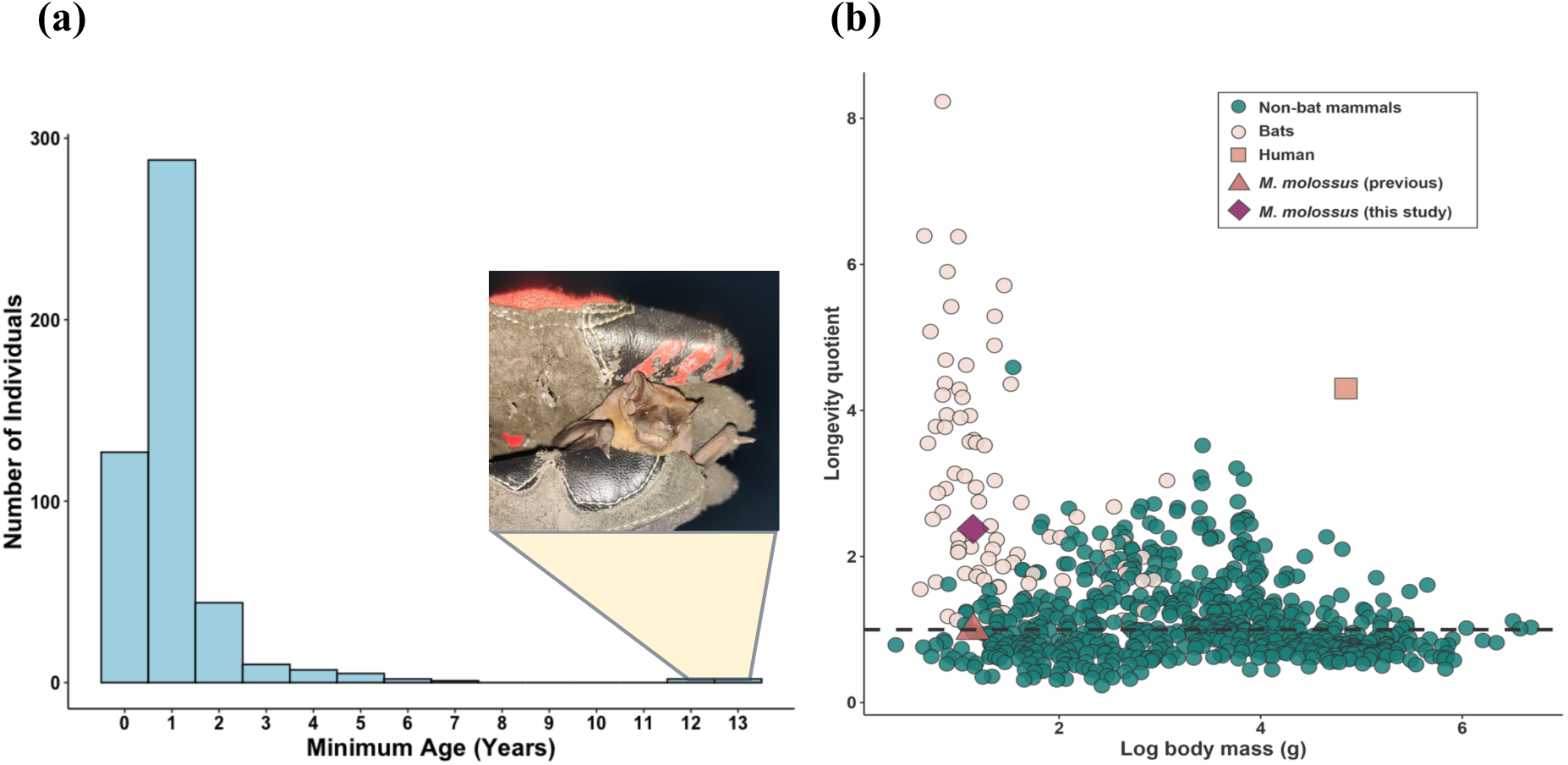
Minimum age distribution and updated longevity quotient for *Molossus molossus*. **(a)** Distribution of minimum age (in years) for *M. molossus* individuals captured between 2021 - 2024. The inset photograph shows Bat 0006C8684C, the oldest known *M. molossus* to date. **(b)** Relationship between longevity quotient (LQ = observed/expected longevity) and log body mass across 829 mammalian species. Each point represents a different species, with non-bat mammals shown as teal circles, bats as pale pink circles, and humans indicated by a light orange square. The previously reported *M. molossus* LQ (Gager et al., 2016) is shown as a red triangle, while the new LQ based on the longevity record discovered in this study is indicated by a purple diamond. The dashed horizontal line represents an LQ of 1, indicating expected longevity based on body mass. Species above this line live longer than predicted for their size. Modified from Foley et al., (2018).

### Telomere length variation with age and sex in *M. molossus*

Accounting for samples which failed DNA extractions, qPCR assay or lacked complete metadata, a total of 466 rTL measurements were calculated for *M. molossus*. Individuals ranged in age from 0 to 13+ years, and repeated sampling resulted in 351 uniquely identified individuals.

We first tested whether telomere length varied with age using a linear mixed effects model including age as a fixed effect and individual identity, assay plate, site, and year as random intercepts (Model 1). This model indicated a significant negative relationship between age and rTL, with telomere length declining with increasing age (β = −0.084 ± 0.033 SE, p = 0.014; Table 1, Figure 2). However, this effect was strongly influenced by the inclusion of the single oldest individual, which contributed three observations at the upper extreme of the age distribution. When this individual was excluded (Model 2, n = 463), the magnitude of the age effect was reduced and no longer statistically significant (β = −0.062 ± 0.048 SE, p = 0.188; Table 1).

**Figure 2:**
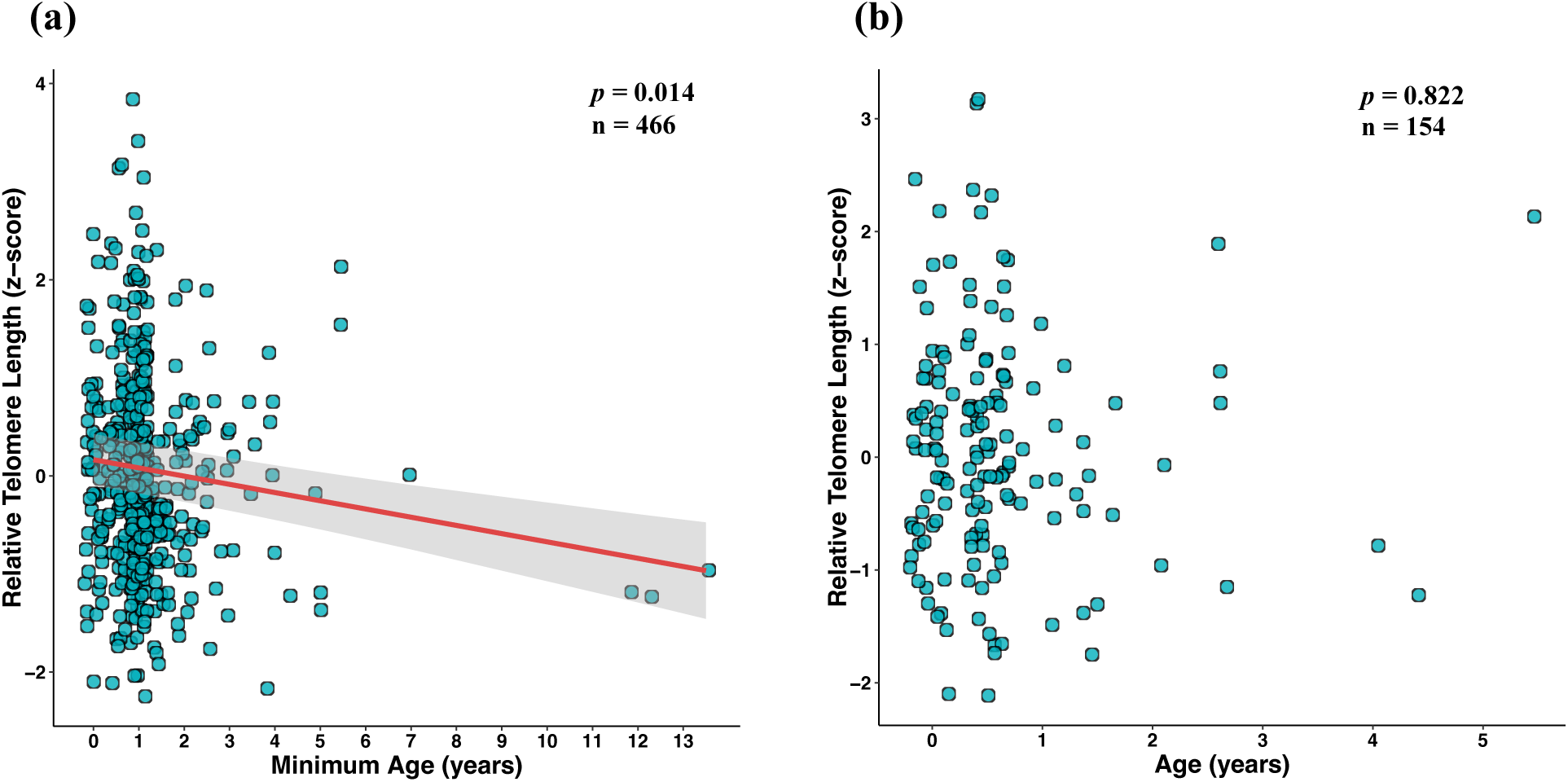
Relationship between age (years) and relative telomere length (rTL) in *Molossus molossus*. **(a)** Full dataset including the oldest individual (0006C8684C; n = 466). The red line represents the linear regression fit suggesting a significant negative relationship between age and rTL (Model 1, p = 0.014). **(b)** Known aged individuals only (n = 154), showing no significant relationship between age and rTL (Model 3, p = 0.822).

**Table 1:**
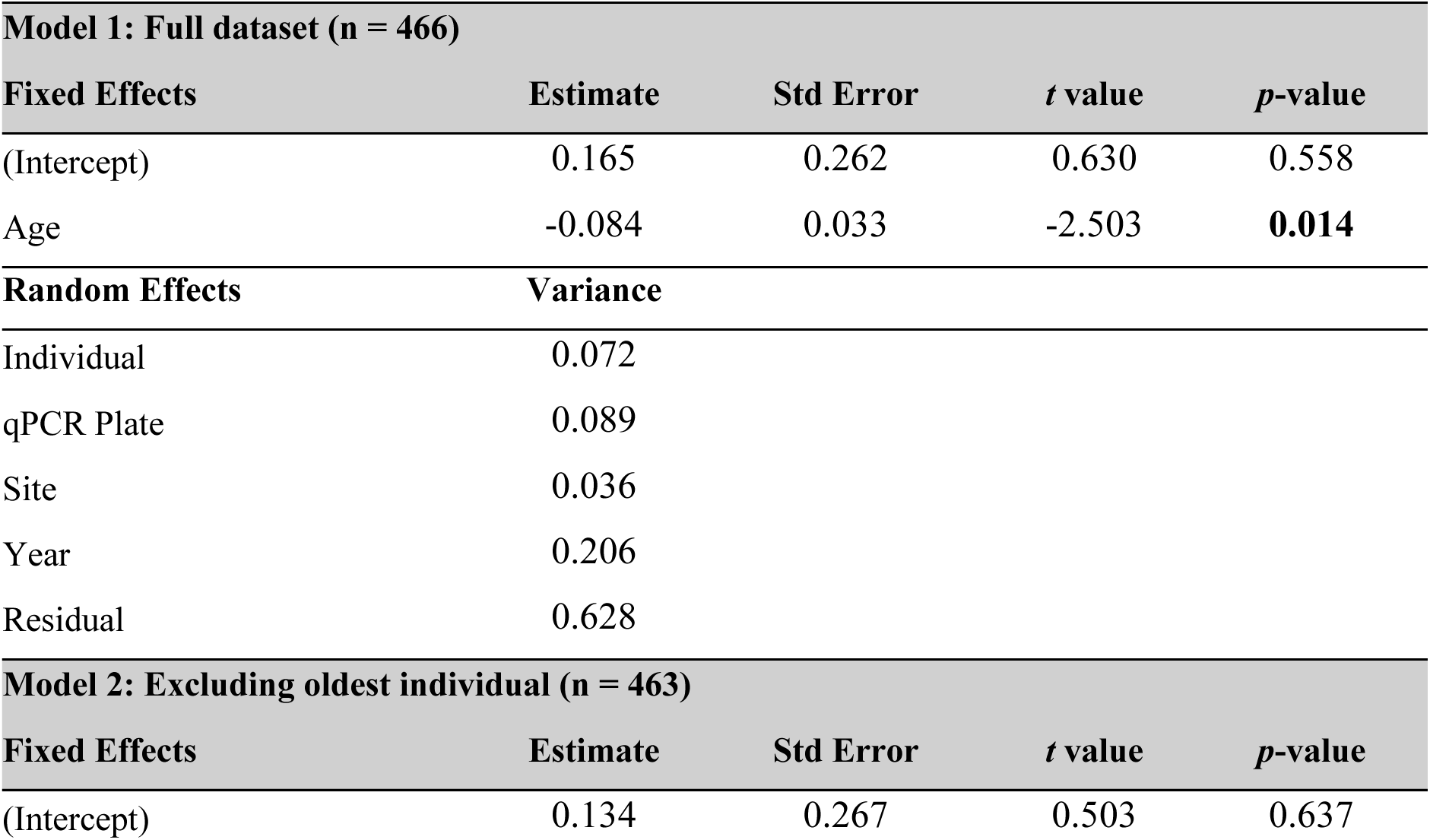
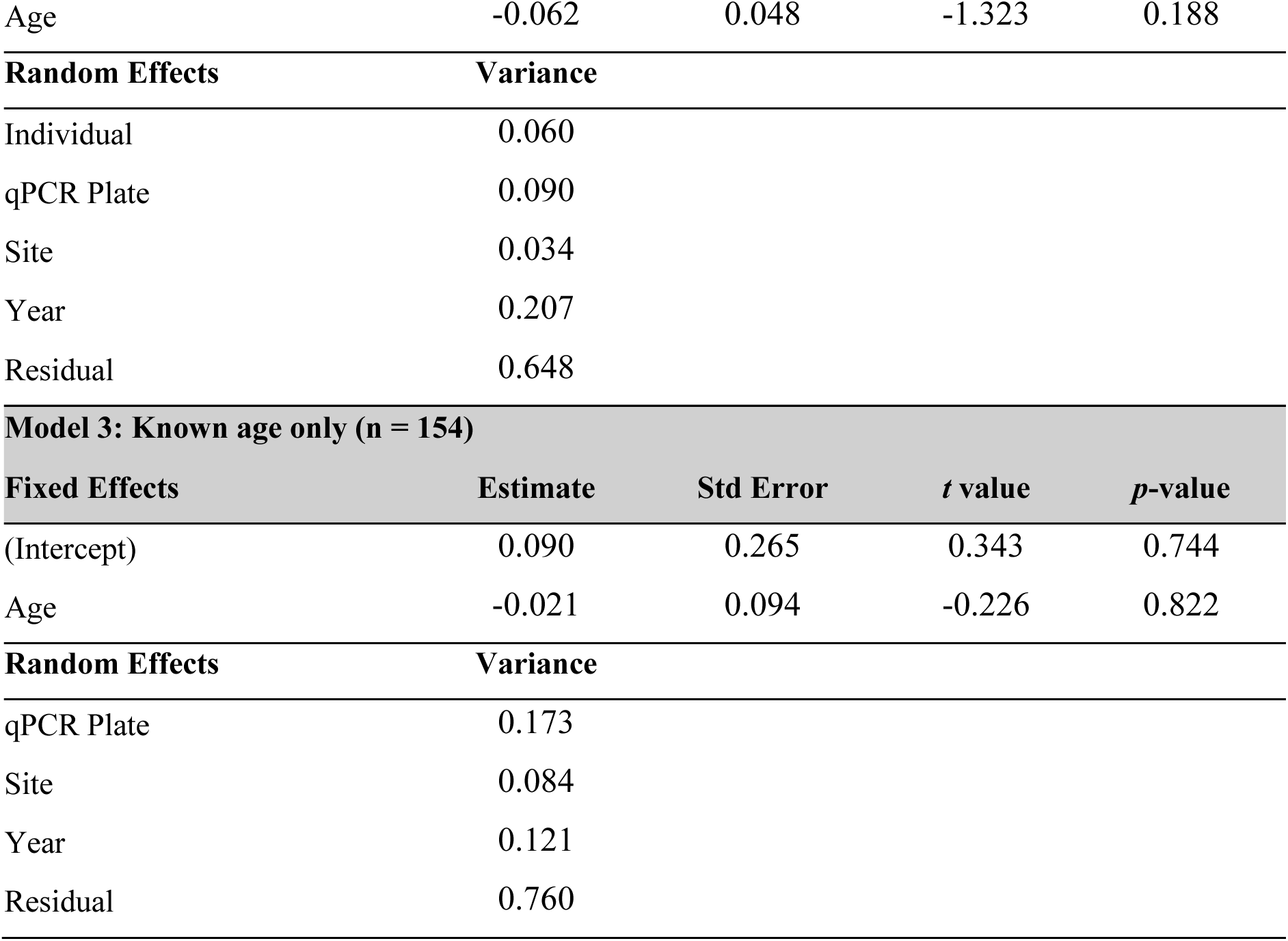
Results of Linear Mixed-Effects Models examining the relationship between age and relative telomere length in *Molossus molossus.* Results shown for three models: the full model including the oldest bat (0006C8684C), a model excluding 0006C8684C and a separate model including only individuals of known age. Models include random intercepts for Individual identity, qPCR plate, capture Site and Year. Significant values in bold p < 0.05.

To evaluate whether this sensitivity reflected disproportionate statistical influence, we fitted robust linear mixed-effects models that down-weight influential observations. In the full dataset, the robust model supported a negative association between age and rTL (β = −0.083; 95% CI: −0.142 to −0.032; Table S2). However, exclusion of the oldest individual again resulted in loss of statistical support for an age effect (β = −0.065; 95% CI: −0.150 to 0.011; Table S2). Thus, statistical evidence for age related telomere shortening was contingent on inclusion of this single extreme age individual, and this individual was excluded from subsequent analyses.

A separate linear mixed effects analysis including only individuals of exact known age (Model 3, n = 154) did not detect a significant relationship between age and rTL (β = −0.021 ± 0.094 SE, p = 0.822; Table 1, Figure 2).

We next examined whether telomere length differed between sexes using a linear mixed effects model including age and sex as fixed effects and individual identity, qPCR plate, site, and year as random intercepts (Model 4). Male *M. molossus* exhibited slightly longer telomeres than females, although this effect was not statistically significant in the standard model (β = 0.165 ± 0.087 SE, p = 0.057; Table 2, Figure 3). To test whether telomere attrition with age differed between sexes, we fitted an interaction between age and sex (Model 5). The interaction term was not statistically supported (β = −0.185 ± 0.131 SE, p = 0.159; Table 2), indicating no evidence for sex specific rates of telomere shortening. Together, these results suggest that males and females follow broadly parallel age related telomere trajectories, with any sex differences primarily reflecting differences in mean telomere length rather than ageing dynamics.

**Figure 3:**
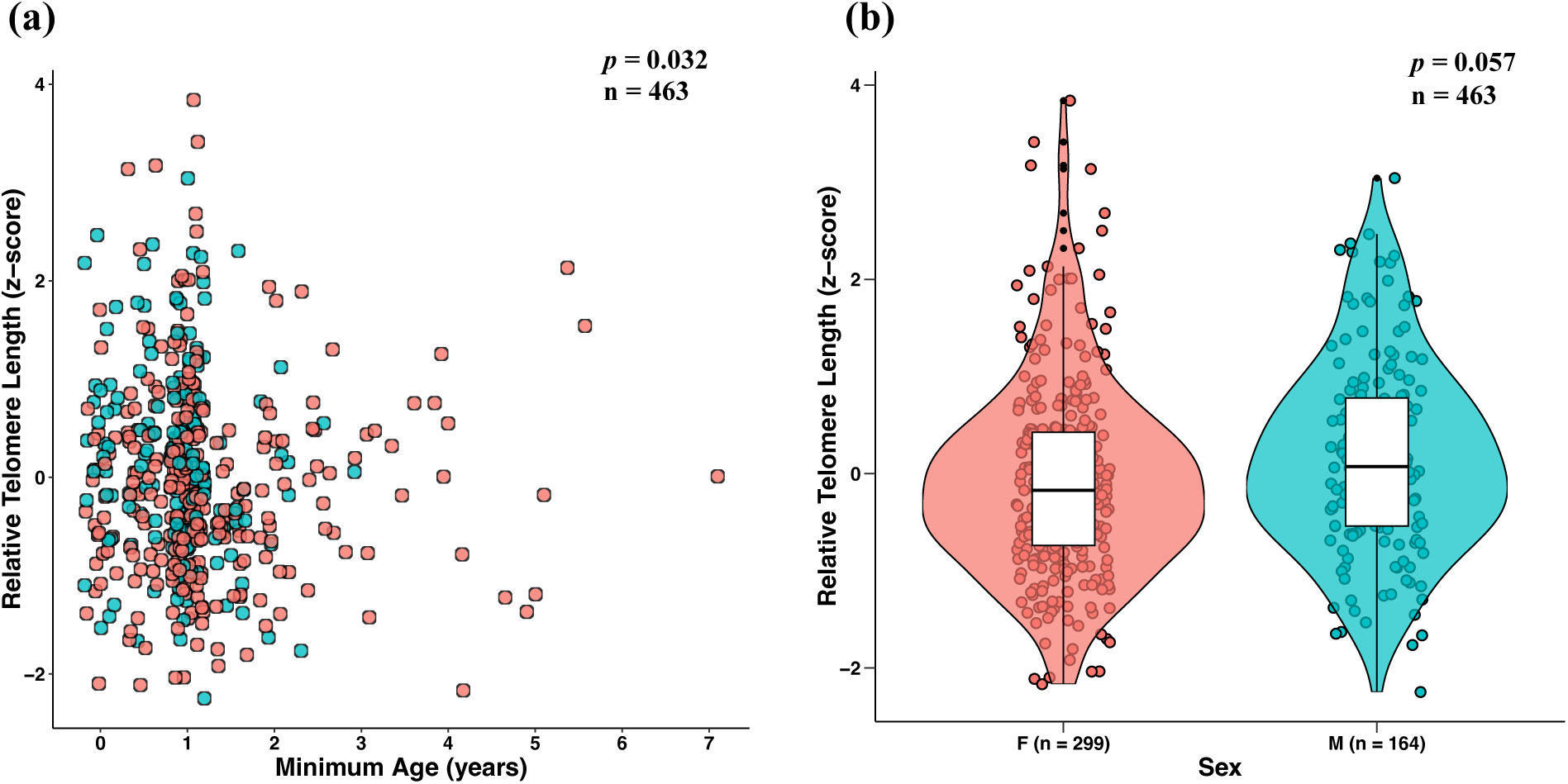
Relationship between age and sex on relative telomere length (rTL) in *Molossus molossus.* **(a)** Relationship between minimum known age (years) and rTL (z-scores). Points represent individual measurements (n = 463), coloured by sex: female (red, n = 299) and male (blue, n = 164). No significant relationship between age and rTL found (Model 4, p = 0.032). **(b)** Violin plot showing the distribution of rTL by sex. Females (F) are shown in red, while males (M) are shown in blue. The boxplot within each violin represents the median and interquartile range. No significant relationship between sex and rTL found (Model 4, p = 0.057).

**Table 2:**
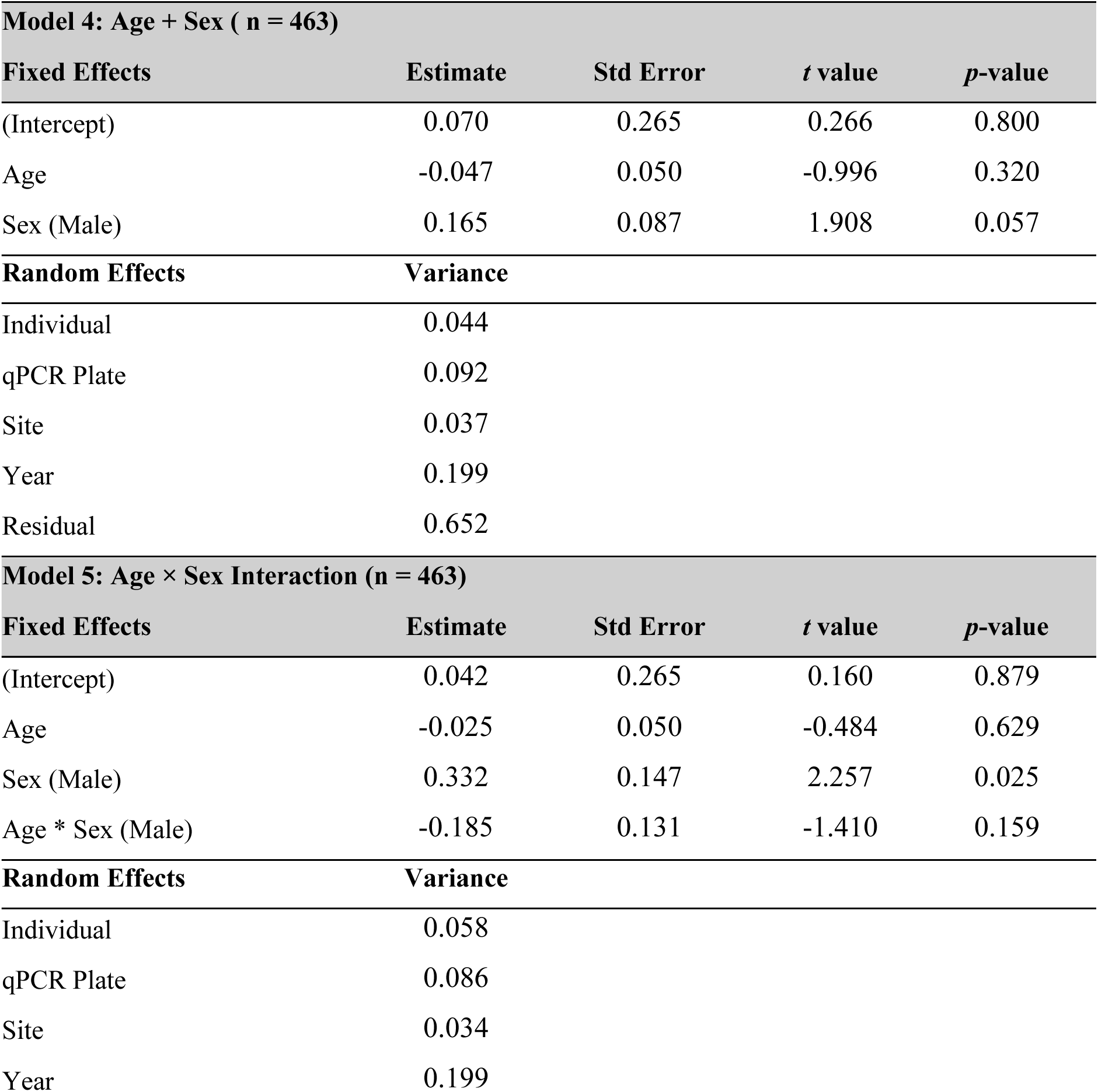
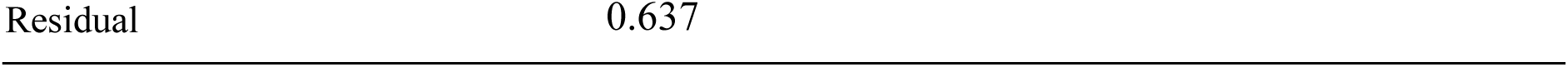
Results of Linear Mixed-Effects Models examining the effects of age and sex on relative telomere length in *Molossus molossus.* Results shown for two models: a model including age and sex as fixed effects and the age-sex interaction model. Age represents the minimum known age in years. Models include random intercepts for Individual identity, qPCR plate, capture Site and Year. Significant values in bold p < 0.05.

Robust versions of the sex models yielded a consistently positive estimate for the effect of male sex on mean rTL (β = 0.340; 95% CI: 0.099 to 0.604; Table S2). In contrast, confidence intervals for the age × sex interaction overlapped zero (β = −0.166; 95% CI: −0.410 to 0.041; Table S2), corroborating the absence of sex specific telomere attrition.

To assess whether observed sex differences could be explained by unequal age distributions, we performed an age-matched subsampling procedure. Across 10,000 iterations in which males were matched to females of identical age, males generally exhibited higher rTL than females. Positive sex-effect estimates were obtained in the majority of subsamples, and the sex effect was statistically significant (p < 0.05) in 73.8% of iterations (Figure S4). Wilcoxon rank-sum tests confirmed that male and female age distributions were successfully matched in all iterations (0% significant tests).

## Discussion

This study represents the first investigation into telomere dynamics in *M. molossus*, a tropical species of bat historically categorised as the shortest-lived bat for which reliable data were available (Tacutu et al., 2018). Through a multi-year mark-recapture study in Gamboa, Panama, we report a new longevity record for *M. molossus*, with an individual female surviving to at least 13 years, significantly exceeding the previously reported maximum lifespan of 5.6 years (Gager et al., 2016). This earlier estimate placed *M. molossus* as a stark outlier compared to temperate-zone bats such as *M. brandtii* (41 years, LQ 8.23), *M. myotis* (37 years, LQ 5.71) and *R. ferrumequinum* (30 years, LQ 4.98) (Gaisler, 2003, Podlutsky et al., 2005, Ransome, 1995). This apparent short lifespan was hypothesised to result from its energetically demanding niche as a non-hibernating, socially foraging insectivorous bat (Gager et al., 2016). However, our findings suggest that the reported short lifespan of this bat likely stemmed from historical under-sampling rather than true biological limits to its longevity. Like many bats, *M. molossus* is challenging to monitor due to its high mobility, skewed population age structure, and the dispersal of juveniles, which limit the chance of recapturing individuals. Long-term mark-recapture studies are logistically difficult to maintain, especially in tropical systems, yet they remain the only reliable method for empirically determining true lifespan in wild bats. The findings here challenge existing assumptions regarding *M. molossus* lifespan and underscore the value of long-term field studies in generating reliable life-history data.

Across vertebrates, telomere length typically shortens with age, including in humans (Müezzinler et al., 2013), chimpanzees (Tackney et al., 2014), reptiles (Ujvari and Madsen, 2009), sheep (Froy et al., 2021) and fish (McLennan et al., 2016). However, bats show marked heterogeneity in telomere ageing patterns. Species within the genus *Myotis* exhibit stable telomere length across adulthood despite exceptional longevity (Foley et al., 2018; Foley et al., 2020; Ineson et al., 2020), whereas *R. ferrumequinum* demonstrates measurable age-related telomere shortening despite comparable longevity to *Myotis* species (Power et al., 2023). In *M. molossus*, age-related telomere dynamics were sensitive to representation at the upper extreme of the age distribution. Inclusion of the single oldest individual produced a statistically supported negative association between age and rTL, whereas exclusion of that individual removed statistical support for age-related shortening. Robust mixed-effects models, which down-weight influential observations, yielded similar effect size estimates and likewise indicated that inference depended on inclusion of this extreme-age data point. Given this sensitivity, we adopt a conservative interpretation that there is no clear evidence of progressive telomere shortening across the broader sampled age range. A separate analysis restricted to individuals of exact known age likewise detected no association between age and rTL. Together, these results suggest that detectable telomere attrition in *M. molossus* is not strongly supported under current sampling, although continued representation of advanced age classes will be essential for resolving late-life dynamics.

Recently, sex-specific patterns of telomere dynamics were reported in the tropical bat species, *Phyllostomus hastatus* (Rayner et al., 2025) where females exhibited age-related telomere shortening while males did not. However, Rayner et al. did not fit a sex × age interaction in their telomere models. Instead they relied on sex-specific analyses due to strongly divergent age distributions between males and females. While sex-specific models can reveal within-sex age patterns, they do not formally test whether the rate of telomere attrition differs between sexes. In contrast, we explicitly tested the sex × age interaction in *M. molossus* using mixed-effects and robust models and found no support that males and females differ in the rate of telomere attrition with age. This indicates that males and females follow broadly parallel age-related telomere trajectories in *M. molossus*. Model estimates suggested that males have slightly longer telomeres, with this pattern more apparent in robust models and in age-matched subsampling. However, the effect was not consistently supported by the standard mixed-effects model.

The lack of a sex difference in telomere dynamics in *M. molossus* is notable given that males might be expected to experience greater physiological costs due to reproductive effort, social competition and hormone profiles. Such factors are widely linked to variation in telomere attrition across taxa (Barrett and Richardson, 2011, Olsson et al., 2011). The harem social structure, with frequent replacement of harem males, in *M. molossus* may impose greater physiological costs on males than females (Gager et al., 2016a, McCracken and Gustin, 1991). This likely elevates reproductive competition, social dominance behaviours, or increased energetic expenditure by males to maintain access to females, all of which can elevate oxidative stress and potentially contribute to telomere attrition. Sexual size dimorphism, males being larger, can also increase cumulative cell divisions during growth, a factor associated with shorter telomeres or faster loss (Barrett and Richardson, 2011, Tombak et al., 2024). Despite these expectations, our results instead align with a meta-analysis which indicated that sex differences in adult telomere length are generally weak, inconsistent, or absent in vertebrates (Remot et al. 2020). The authors found no overall sex differences in telomere length and no systematic differences in telomere attrition rates between males and females across 51 species of vertebrates including mammals, birds, fish and reptiles. They concluded that sex effects on telomere length are typically subtle and species-specific, rather than a general feature of vertebrate biology. Thus, the results we observed in *M. molossus* are not unexpected under current sampling.

Age-matched subsampling indicated that males tended to exhibit longer average telomere length than females across most iterations, even when age distributions were perfectly balanced. While this supports the possibility of a small mean sex difference, this approach does not account for repeated measures, random effects, or variance partitioning, and therefore complements rather than supersedes mixed-effects inference. The discrepancy between subsampling results and hierarchical models likely reflects the small magnitude of the sex effect relative to residual and among-individual variation. As noted by Remot et al. (2020), detecting subtle sex differences in telomere length often requires very large sample sizes, particularly when individual-level variation is high. Our low recapture of older males, due to the social structure of *M. molossus* colonies, may therefore limit our ability to detect late-life attrition in that sex. Taken together, our results indicate that any sex difference in rTL in *M. molossus* is not clearly resolved with our current data, but that further longitudinal sampling and expanded age coverage will be essential to fully elucidate sex-specific telomere dynamics.

Ecologically, *M. molossus* exhibits several traits that may influence telomere dynamics. Individuals concentrate foraging into very brief bouts of about 30 - 60 minutes, typically after sunset and before sunrise (Dechmann et al., 2009, Dechmann et al., 2010). This contrasts sharply with most insectivorous bats, which forage for several hours per night and sometimes in multiple peaks (Davidson-Watts and Jones, 2006, Harding et al., 2022, Jones and Morton, 1992). Outside of these short foraging windows, *M. molossus* spends the majority of the day and night roosting, with telemetry studies recording mean flight activity of only ∼37 minutes per 24 hours (Dechmann et al., 2011). This compressed foraging schedule is combined with exceptionally low daily energy expenditure and frequent use of shallow torpor-like states, even at relatively high ambient temperatures (Dechmann et al., 2011, O’Mara et al., 2017). Such declines in heart rate and metabolic rate are expected to reduce oxidative damage, and studies show that torpor and hibernation can promote telomere maintenance or elongation in mammals, including bats (Turbill et al., 2013, Power et al., 2023). Such ecological buffering mechanisms could plausibly contribute to maintaining telomere stability across much of adulthood.

Interpretation of these results must account for the modest recapture rate (27%), which could reflect both true mortality, emigration and in some cases roost desertion. Inaccessible roosting areas, predation, and human disturbance (e.g. roost sealing or destruction) were all observed and likely contributed to missing individuals. Although we sampled individuals up to 13+ years of age, known ‘old-age’ age classes remain rare. Moreover, many individuals were first sampled as adults, such that ages represent minimum known ages rather than exact chronological ages, which likely adds noise and reduces power to detect late-life telomere dynamics. Future application of DNA methylation-based age estimation (epigenetic clocks) could substantially improve age resolution and strengthen inference in longitudinal studies (Horvath & Raj, 2018; Newediuk et al., 2025; Kratofil et al., 2026). Additionally, selective disappearance of individuals with shorter telomeres could bias apparent age trends upward (Salmón et al., 2017). Our current dataset does not contain sufficient longitudinal recapture of older individuals to formally test this possibility, and therefore we cannot exclude the potential influence of selective survival on observed age patterns. Continued longitudinal sampling of older individuals will be essential for resolving these uncertainties.

This study challenges previous assumptions about the lifespan of *M. molossus*, demonstrating that individuals can live over twice as long as previously recorded. Under current sampling, we find no robust evidence of progressive telomere shortening across adulthood and no clear sex-specific divergence in telomere dynamics in *M. molossus*. These findings contribute to growing evidence that bats exhibit diverse, lineage-specific telomere ageing patterns and that exceptional longevity relative to body size does not correspond to a single telomere trajectory. As longitudinal datasets expand and older age classes become better represented, *M. molossus* will remain an informative system for understanding how ecology, lifespan and telomere dynamics interact in wild mammals.

## Acknowledgments

We thank the members of the Gamboa BatLab for all their help throughout our fieldwork. In particular we thank Gregg Cohen for all of his assistance. We also thank Camila Calderón, Guadalupe García, Lesley Barria, Manon Römkens, Robyn Kelly, Stanley West, Yann Le Bris and Vanessa Mata for their help in the field. We gratefully acknowledge the Smithsonian Tropical Research Centre and MiAmbiente for facilitating this work. This project was funded by Science Foundation Future Frontiers (grant no. 19/FFP/6790) and a European Research Council Synergy grant (grant no. 101118919) awarded to E.C.T.

