## Supplementary File for "Defying expectations: Extended lifespan and limited age-related telomere shortening in the tropical bat species, *Molossus molossus*"

### Supplementary methods

#### Telomere qPCR assay

Relative telomere length (rTL) was measured by real-time quantitative PCR (qPCR) as the concentration of telomeric DNA relative to a single copy gene (SCG) that is constant in number and optimised for use in bats (Cawthon, 2002, Foley et al., 2018). qPCR amplifications were carried out using the Applied Biosciences Quantstudio 7 Flex Real-Time PCR. The mammalian brain-derived neurotrophic factor (*BDNF*) was used as the SCG for this study. Telomere and SCG reactions were carried out on separate plates due to differences in optimum annealing temperatures (Foley et al., 2018, Foley et al., 2020, Power et al., 2021). All primer stocks were used at 50μM concentration. Telomere reactions were carried out in 10μl reaction volumes containing 2.5μl of DNA (~2-5ng/μl), 5μl of Takara Sybr Green (Clontech), 0.027μl of Tel1 primer; 0.09μl of Tel2 primer; 0.2μl ROX (Takara Sybr Green, Clontech) and the remaining volume with PCR grade water (Invitrogen). The compositions of Telomere and SCG reactions were identical except for the primers: 0.12μl *BDNF-F1* and 0.2μl *BDNF-R1*. The following primers were used to amplify the telomere and *BDNF* sequences: Telomere forward *tel1*: (5'-GGTTTTTGAGGGTGAGGGTGAGGGTGAGGGTGAGGGT-3') and reverse *tel2*: (5'-TCCCGACTATCCCTATCCCTATCCCTATCCCTATCC-CTA-3'); *BDNF_F1* (5'-AGCTGAGCGTATGTGACAGT-3') and *BDNF_R1* (5'-TGGGATTACACTTGGTCTCGT-3').

Samples were run on either 96-well or 384-well plates, with six technical replicates. Reactions were amplified using the following cycling conditions: *telomeres* – 30 s at 95°C, followed by 25 cycles of 15 s at 95°C, 2 mins at 54°C and a melt curve; *BDNF* – 30 s at 95°C, 40 cycles of 15 s at 95°C and 1 min at 58°C followed by a melt curve. To account for among-plate variation, a "golden sample" calibrator was run in triplicate on each plate. This calibrator was prepared from pooled *M. molossus* wing tissue. The golden sample serves as an internal reference to ensure consistency across plates and allows normalization of relative telomere length (rTL) values. A negative control was also included on each plate to detect contamination. While real-time qPCR telomere assays detect both terminal and interstitial telomeric sequences (ITS), potentially leading to an overestimation of absolute telomere length (Foote et al., 2013), this does not bias our results because all measurements are relative within this study. As qPCR-based telomere measurement compares telomeric DNA to a single-copy gene within the same sample, rTL reflects proportional differences among individuals rather than absolute telomere length (Bauch et al., 2022). Thus, any contribution of ITS is assumed to be consistent across samples and should not affect the analyses of rTL in this study.

Raw data was exported and the software package LinRegPCR v2020 (Ruijter et al., 2009) was used to calculate baseline correction, per well amplification efficiencies and Cq values (the number of cycles the qPCR amplification curve crosses a set fluorescence threshold). Data were normalised using the golden sample calibrator as reference across both telomere and SCG reactions. The average Cq was calculated across the six replicates for each sample for both amplicons. The coefficient of variation (CoV) was calculated across all technical replicates for each sample and reaction. To maximize data quality while retaining a large dataset the following CoV thresholds were applied: 5% for telomere reactions and 2.5% for single-copy gene (SCG) reactions (Foley et al., 2018, Foley et al., 2020). Replicates exceeding these thresholds were either removed or the sample was excluded from further analysis, depending on the extent of variation. rTL was then calculated for each sample, following the method described by Pfaffl (2001):

$$rTL = \frac{(E_{TEL})^{Cq_{TEL(Calibrator)} - Cq_{TEL(Sample)}}}{(E_{SCG})^{Cq_{SCG(Calibrator)} - Cq_{SCG(Sample)}}}$$

Where $E_{TEL}$ and $E_{SCG}$ are the amplicon and plate specific qPCR efficiencies, $Cq_{TEL(Calibrator)}$ and $Cq_{SCG(Calibrator)}$ are the average Cq values for the "golden sample" calibrator for each amplicon and $Cq_{TEL(Sample)}$ and $Cq_{SCG(Sample)}$ are the average Cq values for individual samples for each amplicon respectively.

### Supplementary figures

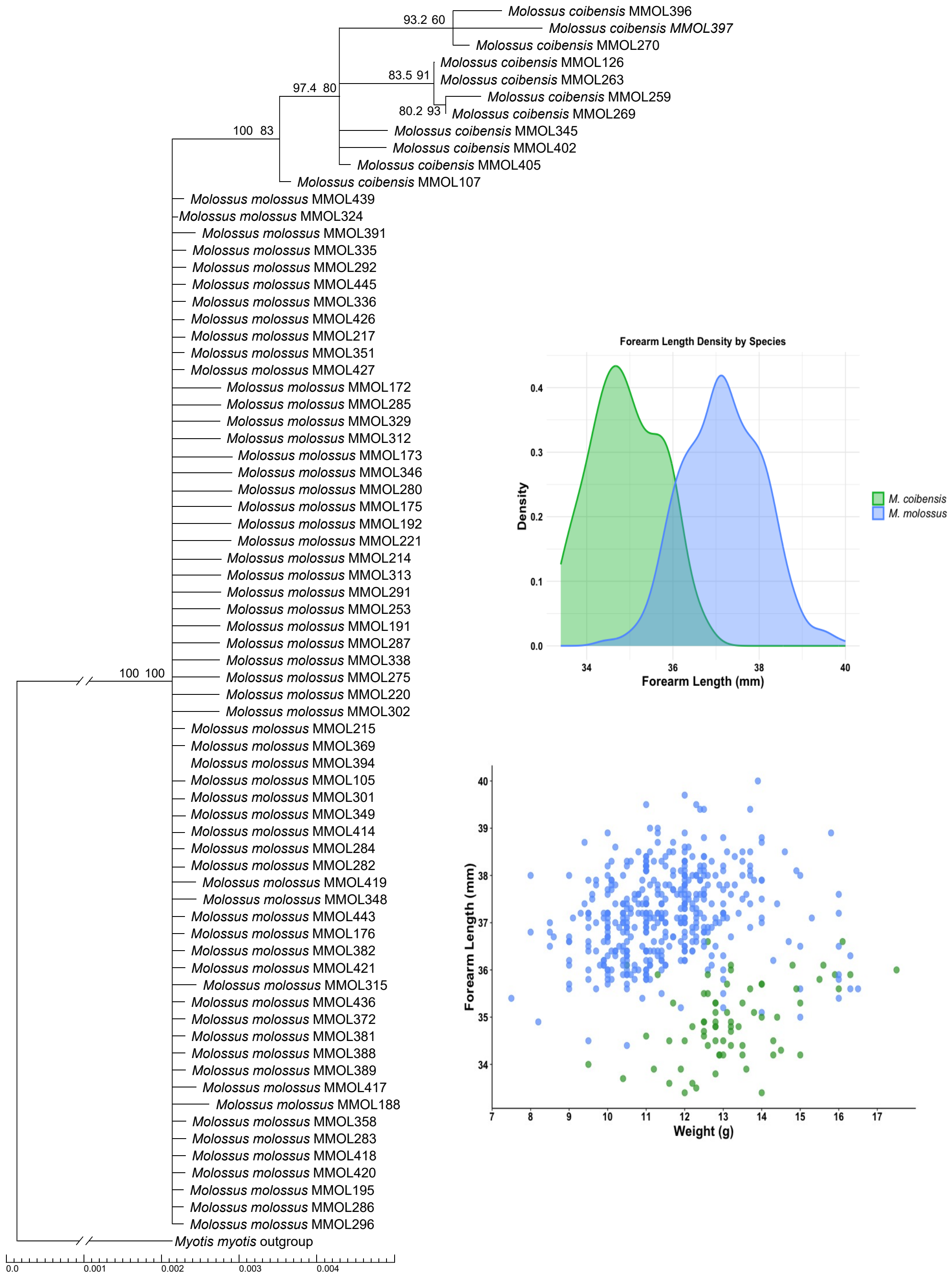


**Figure S1: Genetic and morphological differentiation of *Molossus* species**. **Left**: Maximum likelihood phylogeny based on BRCA1 gene transcripts for 72 molecularly identified individuals. The tree supports species-level clustering, confirming field-based identifications and validating visual classification in the broader dataset. *Myotis myotis* sequence used as an outgroup, sourced from Jebb et al., (2020). Support values are shown on top of the branches as approximate likelihood ratio test (aLRT) first and then ultrafast bootstrap ufBoot. Outgroup branch is shortened as marked by sloped lines; actual length of the outgroup branch is 0.1248. **Top right**: Scatterplot of forearm length (mm) versus weight (g), excluding juveniles and pregnant females. Individuals are coloured by species identity (*M. molossus* = blue, *M. coibensis* = green), showing overlapping but distinguishable morphometric clusters. **Bottom right**: Density plot of forearm length by species, showing clear separation between *M. molossus* and *M. coibensis*, with *M. coibensis* exhibiting generally shorter forearms.

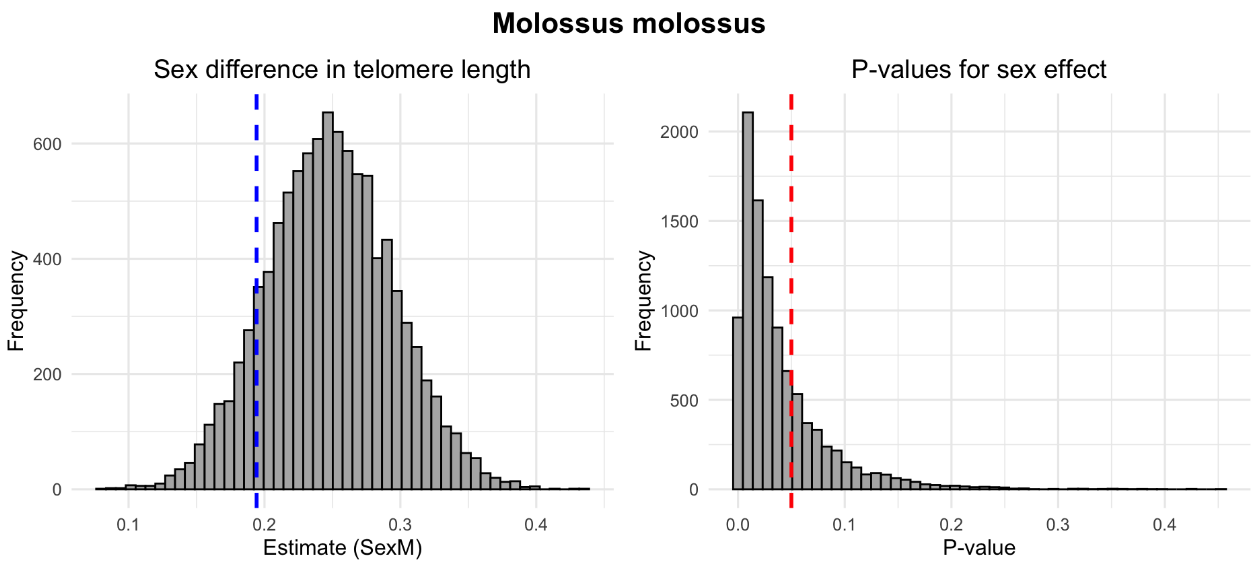


**Figure S2: Sex differences in telomere length after age-matched subsampling for *Molossus molossus*.** Each panel shows results from 10,000 iterations of subsampling in which males were matched to females of the same exact age. In each iteration, a linear model was used to estimate the effect of sex on telomere length. Left panels display the distribution of sex effect estimates (coefficient for males, "SexM") across subsamples. Dashed blue vertical lines indicate the sex effect estimate from the full, unmatched dataset. Right panels show the distribution of p-values for the sex effect across subsamples; red dashed lines mark statistical significance ($p = 0.05$). *M. molossus* displays robust evidence for a sex difference: males consistently show longer telomeres, and the effect was statistically significant in 73.8% of subsamples. Wilcoxon rank-sum tests confirmed that male and female age distributions were successfully matched (0% of iterations showed significant age differences).

#### Supplementary tables

**Table S1: Summary of *Molossus molossus* only captures across roost sites from 2021 to 2024.**

| Roost | 2021 | 2022 | 2023 | 2024 | Females | Males | Juveniles | Sub-Adults | Adults | Roost Total |
|---|---|---|---|---|---|---|---|---|---|---|
| **150** | 19 | 51 | 51 | 52 | 111 | 62 | 21 | 24 | 128 | 173 |
| **152** | 1 | 17 | 30 | 0 | 37 | 11 | 6 | 9 | 33 | 48 |
| **168** | 0 | 0 | 0 | 11 | 7 | 4 | 2 | 0 | 9 | 11 |
| **251a** | 0 | 10 | 0 | 0 | 8 | 2 | 1 | 3 | 6 | 10 |
| **267** | 14 | 25 | 20 | 13 | 57 | 15 | 7 | 13 | 52 | 72 |
| **269** | 9 | 18 | 8 | 0 | 29 | 6 | 5 | 4 | 26 | 35 |
| **275** | 0 | 0 | 8 | 4 | 6 | 6 | 1 | 1 | 10 | 12 |
| **Church** | 66 | 29 | 16 | 13 | 58 | 65 | 9 | 18 | 96 | 124 |
| **School** | 0 | 0 | 7 | 0 | 3 | 4 | 2 | 1 | 4 | 7 |
| **Sub Total** | **109** | **150** | **140** | **93** | **316** | **175** | **54** | **73** | **364** | **492** |

**Table S2: Robust linear mixed-effects models testing the effects of age and sex on relative telomere length in *Molossus molossus*.** Models include: **Model 1**, full dataset including the oldest individual (n = 466); **Model 2**, excluding the oldest individual (n = 463); **Model 3**, excluding the oldest individual and including sex as a fixed effect; and **Model 4**, including an age × sex interaction. Parametric bootstrap confidence intervals were estimated using 10,000 simulations. Random intercepts were included for assay plate, year, and site.

| **Robust Model 1: Age only (full dataset; n = 466)** | | | |
|---|---|---|---|
| **Fixed Effects** | **Estimate** | **2.5% CI** | **97.5% CI** |
| (Intercept) | 0.135 | -0.044 | 0.264 |
| Age | -0.083 | -0.142 | -0.032 |

| **Robust Model 2: Age only (oldest individual excluded; n = 463)** | | | |
|---|---|---|---|
| **Fixed Effects** | **Estimate** | **2.5% CI** | **97.5% CI** |
| (Intercept) | 0.115 | 0.011 | 0.264 |
| Age | -0.065 | -0.150 | 0.011 |

| **Robust Model 3: Age + Sex (oldest individual excluded; n = 463)** | | | |
|---|---|---|---|
| **Fixed Effects** | **Estimate** | **2.5% CI** | **97.5% CI** |
| (Intercept) | 0.028 | -0.088 | 0.202 |
| Age | -0.047 | -0.134 | 0.029 |
| Sex | 0.187 | 0.030 | 0.328 |

| **Robust Model 4: Age x Sex (oldest individual excluded; n = 463)** | | | |
|---|---|---|---|
| **Fixed Effects** | **Estimate** | **2.5% CI** | **97.5% CI** |
| (Intercept) | 0.004 | -0.123 | 0.173 |
| Age | -0.025 | -0.114 | 0.058 |
| Sex | 0.340 | 0.099 | 0.604 |
| Age * Sex | -0.166 | -0.410 | 0.041 |
